# Aerotolerant methanogens use seaweed and seagrass metabolites to drive marine methane emissions

**DOI:** 10.1101/2024.10.14.618369

**Authors:** N. Hall, W. W. Wong, R. Lappan, F. Ricci, R. N. Glud, S. Kawaichi, A-E. Rotaru, C. Greening, P. L. M. Cook

## Abstract

Methanogenesis is classically thought to be limited to strictly anoxic environments. While oxygenated oceans are a known methane source, it is argued that methanogenesis is driven by methylphosphonate-degrading bacteria or potentially is associated to zooplankton gut microbiomes rather than by methanogenic archaea. Here we show through *in situ* monitoring and *ex situ* manipulations that methane is rapidly produced by archaea in frequently oxygenated sandy sediments. By combining biogeochemical, metagenomic, and culture-based experiments, we show this activity is driven by aerotolerant methylotrophic methanogens (*Methanococcoides* spp.) broadly distributed in surface layers of sandy sediments, providing evidence of a hidden process contributing to marine methane emissions. Moreover, we show that methane emissions are driven by methylated seaweed and seagrass metabolites, revealing an unexpected feedback loop between eutrophication-driven algal blooms and greenhouse gas emissions.

## Main Text

Methanogens are typically considered to be strict anaerobes with high sensitivity to oxygen, and therefore restricted to stable anoxic environments (*1, 2*). While some methanogens have been shown to be able to survive periods of oxygen exposure (*3*), active methanogenesis in the environment typically recovers only over timescales of weeks to months (*4–6*).

Of total marine emissions, the contribution of near-shore shallow coastal areas is both the largest and the most uncertain and is estimated to constitute around 75% of marine methane emissions globally, offsetting much of the CO2 drawdown of these highly productive ecosystems (*7*). While the methane emissions of mangroves, salt marshes and other vegetated coastal environments have been actively studied (*8–12*), permeable (sandy) coasts have been largely overlooked, despite covering 50% of the world’s continental margins (*13*).

Methane supersaturation is frequently observed in near-shore waters overlying permeable sediments, and has generally been explained by input of methane-rich groundwater or riverine water, or seepage of methane from below the sulfate-methane transition zone (*14*– *16*). It has also been proposed that excess methane in these zones could be produced by aerobic bacteria during methylphosphonate degradation (*17, 18*) or through phytoplanktonic photosynthesis processes (*19–22*). Archaeal methanogenesis has been disregarded as a significant contributor given coastal permeable sediments are characterized by sudden and unpredictable changes in redox conditions and high sulfate concentrations (*23, 24*). These characteristics promote the dominance of metabolically flexible facultative anaerobic bacteria (*25*) and are thought to exclude methanogenic archaea.

Here we demonstrate methylotrophic methanogenesis is active in surface sediments under short-term anoxia and is stimulated by seaweed and seagrass (collectively macrophyte) metabolites. By pairing isolation of two strains of *Methanococcoides* sp. with metagenomic profiling, we show that archaeal methylotrophic methanogens are widespread and active in sandy sediments at sites of both Australia and Europe.

### High methane levels occur in near-shore surface waters due to sedimentary production

Methane concentrations measured in near-shore surface waters over beaches around Port Phillip Bay and Westernport Bay (Australia) and Avernakø (Denmark) are consistently oversaturated with respect to the atmosphere. The extent of saturation was shown to vary over four orders of magnitude, with values ranging from 300% to 160,000% saturated with respect to the atmosphere (Fig. 1A). No relationship was found between methane and radon concentrations (Fig. 1B), indicating methane was produced in either permeable sediments or surface waters rather than through groundwater seepage, contrasting with similar previously described cases such as in the Gulf of Mexico and South Sea of Korea (*26, 27*).

**Figure 1.**
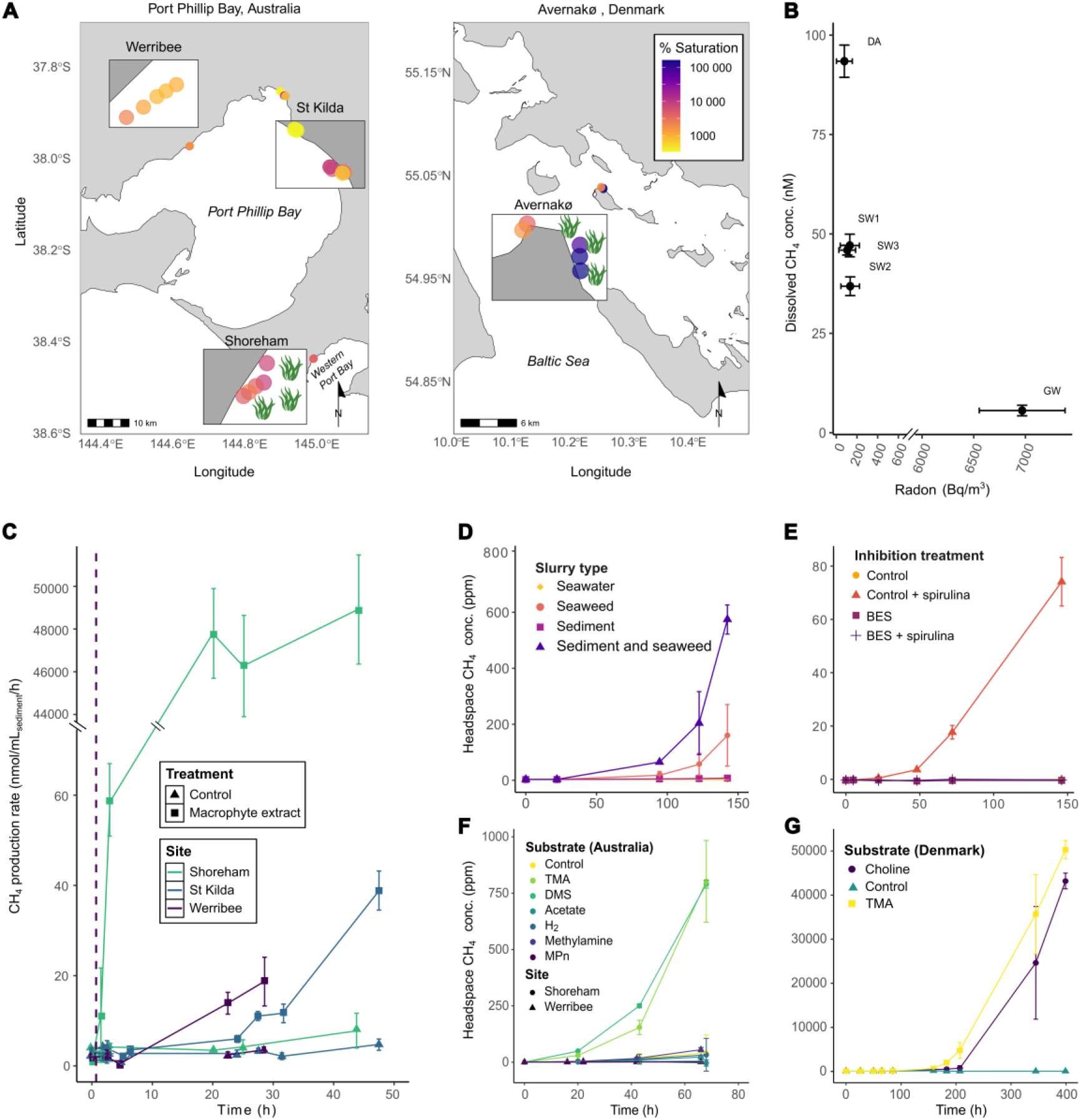
(**A**) Map of *in situ* surface water methane supersaturation at Australian and Danish sites. Green seaweed symbols indicate sites with high macrophyte accumulation. For detailed site information see table S1. (**B)** Methane (nM) versus radon (Bq/m^3^) concentrations in groundwater (GW), surface water clear of drift algae (SW) and surface water from within a drift algal mat (DA) at Werribee. Error bars indicate standard deviation of triplicate samples (methane) or error given by ^222^Radon analyzer. (**C**) Flow-through reactor (FTR) experiments show methane production after onset of anoxia and macrophyte extract addition (dashed line) in surface sediments (0-5 cm) from a site with macrophyte accumulation (Shoreham) and sites without accumulation (Werribee and St Kilda). Error bars represent standard deviation from mean of four independent FTRs. Note broken y axis. (**D**) Methane production in slurry incubations of seawater, sediment and seaweed (brown drift algae) from Werribee. (**E**) Methane production in Werribee sediment slurry incubations with spirulina additions with and without specific archaeal methanogenesis inhibitor BES (20 mM). (**F**) Methane production in sediment slurry incubations with addition of 100 μM concentrations of trimethylamine (TMA), dimethyl sulfide (DMS), acetate, hydrogen (340 ± 20 ppm), methylamine and methylphosphonate (MPn) in sediments from Shoreham and Werribee. (**G**) Methane production in sediment slurry incubations with addition of 100 μM concentrations of trimethylamine (TMA) and choline in sediments from Avernakø, Denmark. All slurries were prepared with surface sediment (0-5 cm) and argon-purged at the start (time = 0 h). Error bars of all slurry experiments represent standard deviation from mean of three independent slurries.

While methane was always supersaturated with respect to the atmosphere, higher methane concentrations were observed over local (meters to tens of meters) scales adjacent to, or within, seaweed and seagrass mats at both Werribee (Fig. 1B) and Avernakø (Fig. 1A), indicating that this biomass enhances methane production in the environment. To determine whether methane production specifically occurs in the sediment, water column, or on the surface of the seaweed or seagrass biomass, rates of methane production were compared between slurry incubations with combinations of sediment, seawater, and seaweed. The combination of sediment and seaweed stimulated the highest rates of methane production, followed by seaweed only (Fig. 1D), whereas no methane production was detectable in seawater-only controls or sediment only. This indicates that microbes responsible for methane production are primarily located in sediment, but benefit from substrates derived from seaweed or seagrass.

### Methylotrophic methanogenesis driven by plant metabolites dominates emissions

We initially hypothesized that bacteria or microalgae, rather than methanogenic archaea, may be responsible for methane production in permeable sediments because frequent oxygenation of the surface sand would inhibit archaeal methanogenesis (*28*). 2-bromoethane sulfonate (BES) was used as a targeted inhibitor of archaeal methanogenesis through its inhibition of methyl-CoM reductase, the terminal enzyme in all known pathways of archaeal methanogenesis (*29, 30*). BES addition completely inhibited methane production in our slurries, indicating that the only methane-producing microbes were methanogenic archaea (Fig. 1F). Methylphosphonates did not stimulate methane production in either slurries (Fig. 1G) or oxic or anoxic flow through reactors (FTRs) (Fig. S3), further ruling out a bacterial source and showing that the pathway of methane production in this environment is different to that which has been used previously to explain the oceanic methane paradox in marine surface waters (i.e. methylphosphonate degradation in the water column) (*31–33*).

We investigated acetoclastic, hydrogenotrophic, and methylotrophic methanogenesis pathways using targeted substrate addition (Fig 1F) and found that methylotrophic methanogenesis predominated. This is likely due to the fact that many organisms such as sulfate and nitrate reducers outcompete methanogens for acetate and hydrogen, but not for methylated substrates, allowing methylotrophic methanogenesis to occur in unreduced environments with abundant alternative electron acceptors (*34, 35*). Additionally, surface sediments are often exposed to high fluxes of labile methylated compounds and may therefore be adapted to quickly utilize these substrates as described in seagrass meadows (*36*). Methylotrophic methanogenesis was similarly stimulated by trimethylamine (TMA) and dimethyl sulfide (DMS) (Fig. 1F); these are abundant compounds in coastal regions and are formed through the breakdown of widely occurring macrophyte osmolytes such as glycine betaine, choline, and dimethyl sulfoniopropionate (DMSP) (*37, 38*).

The rate of potential methane production from sediment at sites with different levels of macrophyte accumulation was examined with FTR experiments to investigate the role of community composition and priming effects. FTRs more realistically emulate the advective flow which is caused by wave and tidal pumping in surface permeable sediments (*39, 40*). Remarkably, methanogenesis started within ∼20 hours of the transition to anoxia and addition of macrophyte extract in FTRs from all sites (Fig. 1C), which is much faster than previously reported recovery times of weeks to months (*4–6*), regardless of site. However, the rate of methane production at the site with the most macrophyte accumulation (Shoreham) was both faster (within 1.5 hours) and four orders of magnitude higher (at 20 hours) than the two sites with less accumulation. This indicates that permeable sediments may generally harbor the latent potential for methanogenesis, but supports our hypothesis that macrophyte accumulation causes changes in abundance and/or activity of methanogens, resulting in higher methane production rates.

While methane production rates are difficult to compare between different sediment types due to the differences in advective and diffusive flux, some comparisons can be made between our experimental rates to place them in the context of more well-studied environmental fluxes. If integrated conservatively over a 0.5 cm permeable sediment depth as the depth of advective penetration of high-substrate surface water in the intertidal zone (*41*), the maximum methane production rate at Shoreham is approximately 3.8 g/m^2^/h, nearly three orders of magnitude higher than flux rates recently reported for tropical wetlands (*42, 43*). The CH4:CO2 carbon remineralization ratio reached 1:9 after 44 hours (Fig. S4), exceeding the typical ratio for most types of wetland sediment and indicating that methanogenesis may be a quantitatively important carbon remineralization path in permeable sediments (*44*). These findings highlight not only the ability of methanogenic archaea to adapt highly dynamic environments and recover extremely quickly after oxygen exposure, but also that their methane production rates can be comparable to, or even exceed, those of the most active methanogenic environments studied (*44–46*). Together, these field data and inhibitor slurry experiments show that archaeal methanogenesis is not only feasible in surface sandy sediments, but in some cases, appears to be the major source of methane to the overlying water and subsequent emissions. Furthermore, these results reveal a mechanism linking macrophytes to elevated methane concentrations, which could be significant in the context of increasing ocean eutrophication and enhanced macrophyte growth.

### Novel archaeal isolates and sediment metagenomes confirm aerotolerant methanogenic potential

Methanogens were isolated from both Australian and Danish sites to investigate the aerotolerance of methanogens from permeable sediments, independent of potential anaerobic sand grain microniches caused by bacterial oxygen consumption. Two methanogens from the genus *Methanococcoides* (family *Methanosarcinaceae*) were isolated from surface sands (0-5 cm), one from Avernakø, Denmark (strain DA) the second from Shoreham, Australia (strain SH). Based on average nucleotide identity (ANI) comparisons with all representative genomes of *Methanococcoides*, DA was most closely related to *Methanococcoides burtonii* (ANI value 89.3) while SH was most closely related to *Methanococcoides orientis* (ANI value 90.0). Between the two isolates, the ANI value was 80.3. In support of the biogeochemical observations, both methanogens rapidly produced methane in the presence of TMA (Fig. 2A). Moreover, activity of these methanogens resumed immediately following transient (approximately 30 min) exposure to 6-9 mg/L oxygen (Fig 2A).

**Figure 2.**
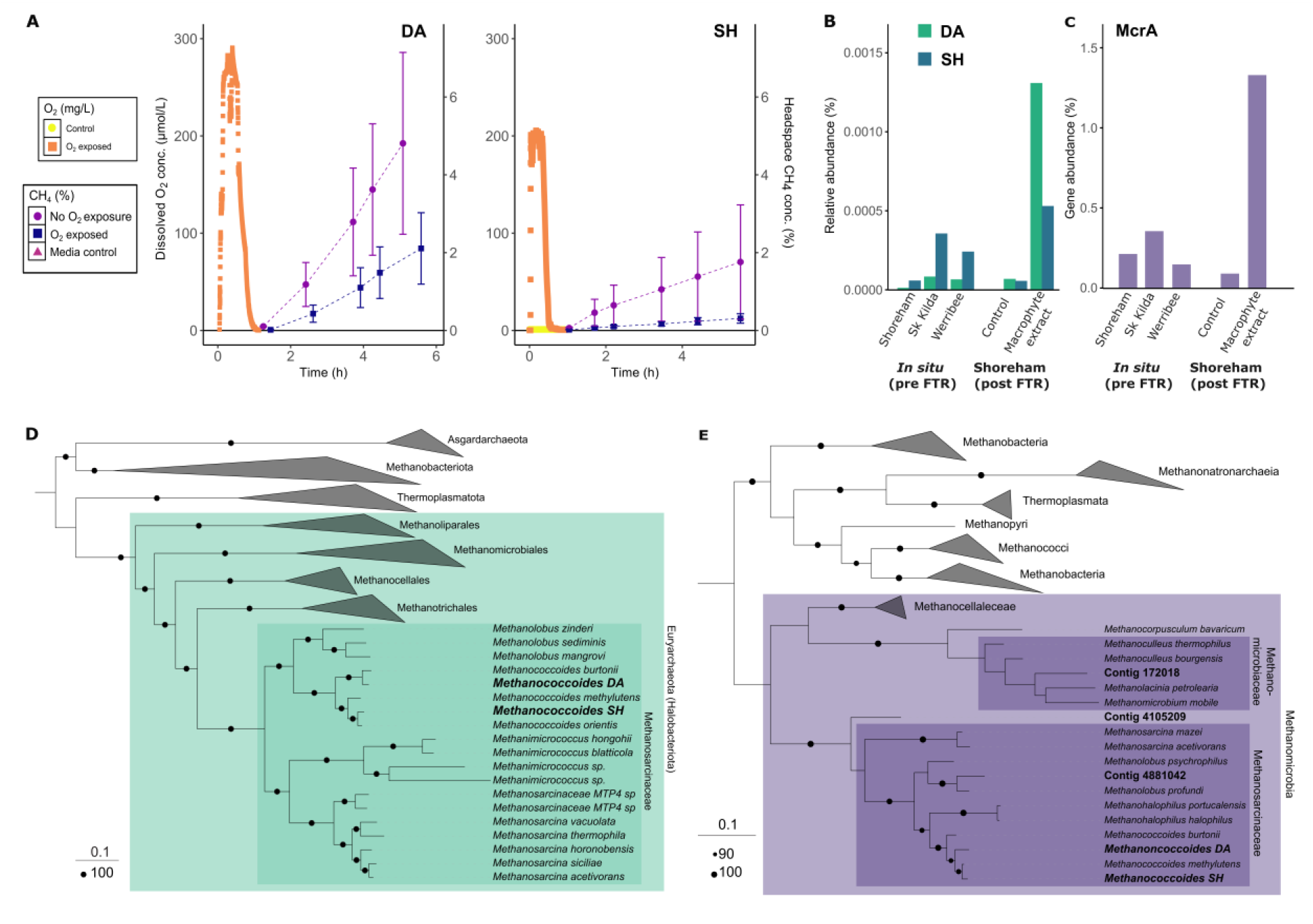
(**A**) Methane production after oxygen exposure in sand isolates *Methanococcoides* sp. DA and SH. (**B**) Read mapping of metagenomic reads to isolate sequences from post-FTR samples (Shoreham) and *in situ* samples from three sites collected at the same time as the FTR sediment samples. (**C**) *mcrA* gene abundance in all metagenome samples. (**D**) Maximum-likelihood phylogenetic genome tree showing both isolates (bold) and reference genomes. Amino acid sequences from the isolated genomes were aligned with reference genomes. Coloured boxes indicate family (dark shade) and phylum (light shade). Legends for tree scale and bootstrap of 100 are reported in the bottom left corner. (**E**) Maximum-likelihood phylogenetic gene tree of *mcrA* sequences, displaying isolates (bold), contigs from metagenomic analyses of in situ and post-FTR sand samples (bold), and reference sequences. Amino acid sequences of isolates and contigs were aligned with reference sequences. Coloured boxes indicate family (dark shade) and class (light shade). Legends for tree scale and bootstrap of 90 and 100 are reported in the bottom left corner. Sequence alignments used to generate these phylogenetic trees are provided and source trees are available in tree file format (see data and materials availability below).

Read mapping of metagenomic reads to the isolate genomes revealed that both strains were present in *in situ* samples from all sites and their abundance increased 9 to 10-fold with the macrophyte extract treatment in FTRs (Fig. 2B). Furthermore, analyses of the genome sequences of both isolates revealed remarkable similarities in their methanogenesis pathways and antioxidant systems, despite being isolated from geographically and climatically distinct locations, suggesting that these traits are important for adaptation in this environmental niche. Both isolates encoded a complete methylotrophic methanogenesis pathway with methyltransferases for tri-, di- and monomethylamines and methanol (*mttABC, mtbABC, mtmABC and mtaABC* respectively). No methylthiol methyltransferases (*mts*) were found, despite DMS equally stimulating methane production to TMA in slurries (*47*). It has been proposed that some methanogens may use *mtt* and *mta* methyltransferases to metabolize DMS (*47*). Both genomes encode various genes involved in oxidative stress protection, including the F420H2-dependent oxidase that detoxifies O2 to water and is unique to methanogens (*48*), as well as those typically associated with aerobic organisms such as superoxide reductase and catalase-peroxidase (*49*). Additionally, both genomes encoded a variety of other proteins with potential antioxidant roles such as rubredoxins (*50*), thioredoxins (*51*), and peroxiredoxins (*3, 52, 53*). This abundance and diversity of genes involved in oxidative stress protection supports the finding of remarkable robustness to, and quick recovery from, oxygen exposure.

Shotgun metagenomics was undertaken to determine if methanogens and their functional genes were present in the permeable sediments. Samples for analysis were collected at the beginning of the FTR experiment for all sites, as well as at the end of the experiment for Shoreham samples. The marker gene for archaeal methanogenesis, *McrA* (encoding methyl-CoM reductase), were found in samples from all sites (Fig. 2C). This gene was encoded by a relatively small proportion of the community (0.094 – 1.33%) in line with previous observations for surface permeable sediments (*25, 54, 55*); however, it’s well-established that methanogens are highly transcriptionally and biogeochemically active even in niches where their abundance is low (*11*). Furthermore, *McrA* gene abundance increased from 0.22% to 1.33% (a six-fold increase) in the Shoreham FTR experiment macrophyte extract treatment but not in the control, indicating growth of methanogens. Most sequences were affiliated with *Methanosarcinaceae* (Fig. 2E), all of which are known to utilize methylated substrates (*35*), giving a genetic basis for the experimental finding that all sites harbor the latent potential for methylotrophic methanogenesis. This suggests a dormant pool of these methanogens exists in sands, with growth stimulated by macrophyte metabolites and anoxic conditions.

### Potential feedback pathways amid ongoing local and global pressures on coastal ecosystems

By integrating *in situ* and *ex situ* data, spanning culture-independent and culture-based experiments, we observe a previously undescribed mechanism for methane production in coastal environments. Methanogens can be adapted to more frequently oxygenated environments than previously thought and sandy coasts broadly harbor the potential to convert macrophyte-derived substrates to methane. These methanogens may help to explain the oceanic methane paradox in marine surface waters of coastal settings, in addition to the previous explanation of methylphosphonate degradation in the water column (*31–33*).

The shallow and turbulent nature of waters overlying coastal permeable sediments, combined with advective transport in the sediments, gives the study additional significance. In deeper waters and cohesive sediments, the balance of methanogenesis and methanotrophy is such that in the bulk of the ocean volume is undersaturated in methane with respect to the atmosphere (*56*). However, in rippled permeable sediments the redox seal is broken, with flow from reduced reaction zones exported directly through ripple peaks or lee sides depending on bedform and flow interactions (*57, 58*). This allows methane produced in shallow anoxic regions to bypass the methane oxidation filter and reach the shallow overlying water where low residence times and high turbulence causes high rates of export to the atmosphere (*14, 59*). Therefore, the contribution of methane production in shallow permeable sediments to total marine methane emissions is likely disproportionately large.

Furthermore, the link between macrophyte biomass and methane emissions has significance in the context of current and predicted ocean changes due to climate change and other anthropogenic impacts. Eutrophication and rising sea temperatures in particular are linked to increased algal growth and the frequency of large algal blooms in coastal zones (*60*– *63*). Here, we have shown that deposition of this excess algal biomass on sandy coasts may result in increasingly large and frequent pulses of methane to the atmosphere and should be accounted for in future marine methane budgets and modelling. As well as unintentional excess macrophyte growth caused by eutrophication, the results of this study further complicate CO2 removal by macrophytes or “blue carbon” as a climate change mitigation strategy (*7*), as enhanced methane emissions may offset much of the CO2 removal by these ecosystems.

## Acknowledgments

The authors would like to thank Rodney Hall, Vera Eate, and Michael Tolj for their technical assistance in sample analysis. Thanks to Anni Glud for experimental help and Isabell Schlangen and Annabell Moser for field assistance.

## Funding

Australian Research Council grant DP210101595 (PLMC, CG, WWW) National Health & Medical Research Council fellowship APP1178715 (CG) Australian Government Research Training Program Scholarship (NH) European Research Council ERC 101045149 (AER, SK)

Danish Research Council DFF 1026-00159 (AER)

Danish National Research Foundation, DNRF145 (RNG, NH)

## Author contributions

Conceptualization: NH, PLMC, CG Methodology: NH, WWW, RL, SK, FR, AER Investigation: NH, RL, FR

Visualization: NH, FR

Funding acquisition: CG, PLMC, WWW, AER Project administration: PLMC, CG, RNG Supervision: CG, PLMC, RNG

Writing – original draft: NH

Writing – review & editing: NH, PLMC, CG, RL, FR, AER, SK, WWW, RNG

## Competing interests

Authors declare that they have no competing interests.

## Data and materials availability

All scripts for bioinformatic analysis, supplementary trees and tree files available at https://github.com/GreeningLab/Sand-methanogen-manuscript Sequence data including metagenomes, isolate genomes, and contigs used for analysis available at NCBI Sequence Read Archive BioProject ID: PRJNA1165813 https://www.ncbi.nlm.nih.gov/bioproject/PRJNA1165813

## Notes

https://github.com/GreeningLab/Sand-methanogen-manuscript

https://www.ncbi.nlm.nih.gov/bioproject/PRJNA1165813

### Competing Interest Statement

The authors have declared no competing interest.

## References and Notes

1. Y. Liu, W. B. Whitman, Metabolic, Phylogenetic, and Ecological Diversity of the Methanogenic Archaea. Annals of the New York Academy of Sciences 1125, 171–189 (2008).

2. T. Hoehler, N. A. Losey, R. P. Gunsalus, M. J. McInerney, “Environmental Constraints that Limit Methanogenesis” in Biogenesis of Hydrocarbons, A. J. M. Stams, D. Sousa, Eds. (Springer International Publishing, Cham, 2018; 10.1007/978-3-319-53114-4_17-1), pp. 1–26.

3. R. Jasso-Chávez, M.G. Santiago-Martínez, E. Lira-Silva, E. Pineda, A. Zepeda-Rodríguez, J. Belmont-Díaz, R. Encalada, E. Saavedra, R. Moreno-Sánchez, Air-Adapted Methanosarcina acetivorans Shows High Methane Production and Develops Resistance against Oxygen Stress. PLoS ONE 10, e0117331 (2015).

4. T. Watanabe, M. Kimura, S. Asakawa, Distinct members of a stable methanogenic archaeal community transcribe mcrA genes under flooded and drained conditions in Japanese paddy field soil. Soil Biology and Biochemistry 41, 276–285 (2009).

5. K. Ma, Y. Lu, Regulation of microbial methane production and oxidation by intermittent drainage in rice field soil. FEMS Microbiology Ecology 75, 446–456 (2011).

6. R. Angel, P. Claus, R. Conrad, Methanogenic archaea are globally ubiquitous in aerated soils and become active under wet anoxic conditions. ISME J 6, 847–862 (2012).

7. F. Roth, E. Broman, X. Sun, S. Bonaglia, F. Nascimento, J. Prytherch, V. Brüchert, M. Lundevall Zara, M. Brunberg, M. C. Geibel, C. Humborg, A. Norkko, Methane emissions offset atmospheric carbon dioxide uptake in coastal macroalgae, mixed vegetation and sediment ecosystems. Nat Commun 14, 42 (2023).

8. S. J. E. Krause, T. Treude, Deciphering cryptic methane cycling: Coupling of methylotrophic methanogenesis and anaerobic oxidation of methane in hypersaline coastal wetland sediment. Geochimica et Cosmochimica Acta 302, 160–174 (2021).

9. J. Yuan, D. Liu, Y. Ji, J. Xiang, Y. Lin, M. Wu, W. Ding, Spartina alterniflora invasion drastically increases methane production potential by shifting methanogenesis from hydrogenotrophic to methylotrophic pathway in a coastal marsh. Journal of Ecology 107, 2436– 2450 (2019).

10. H. J. Jones, E. Kröber, J. Stephenson, M. A. Mausz, E. Jameson, A. Millard, K. J. Purdy, Y. Chen, A new family of uncultivated bacteria involved in methanogenesis from the ubiquitous osmolyte glycine betaine in coastal saltmarsh sediments. Microbiome 7, 120 (2019).

11. M. Cai, X. Yin, X. Tang, C. Zhang, Q. Zheng, M. Li, Metatranscriptomics reveals different features of methanogenic archaea among global vegetated coastal ecosystems. Science of The Total Environment 802, 149848 (2022).

12. B. D. Eyre, N. Camillini, R. N. Glud, J. A. Rosentreter, The climate benefit of seagrass blue carbon is reduced by methane fluxes and enhanced by nitrous oxide fluxes. Commun Earth Environ 4, 1–9 (2023).

13. S. J. Hall, The continental shelf benthic ecosystem: current status, agents for change and future prospects. Environmental Conservation 29, 350–374 (2002).

14. T. Weber, N. A. Wiseman, A. Kock, Global ocean methane emissions dominated by shallow coastal waters. Nat Commun 10, 4584 (2019).

15. M. J. Reading, D. T. Maher, I. R. Santos, L. C. Jeffrey, T. J. Cyronak, A. McMahon, D. R. Tait, Spatial Distribution of CO2, CH4, and N2O in the Great Barrier Reef Revealed Through High Resolution Sampling and Isotopic Analysis. Geophysical Research Letters 48, e2021GL092534 (2021).

16. C. S. Wu, H. Røy, D. de Beer, Methanogenesis in sediments of an intertidal sand flat in the Wadden Sea. Estuarine, Coastal and Shelf Science 164, 39–45 (2015).

17. M. Kanwischer, T. Klintzsch, O. Schmale, Stable Isotope Approach to Assess the Production and Consumption of Methylphosphonate and Its Contribution to Oxic Methane Formation in Surface Waters. Environ Sci Technol 57, 15904–15913 (2023).

18. J. N. von Arx, A. T. Kidane, M. Philippi, W. Mohr, G. Lavik, S. Schorn, M. M. M. Kuypers, J. Milucka, Methylphosphonate-driven methane formation and its link to primary production in the oligotrophic North Atlantic. Nat Commun 14, 6529 (2023).

19. Q. Wang, A. Alowaifeer, P. Kerner, N. Balasubramanian, A. Patterson, W. Christian, A. Tarver, J. E. Dore, R. Hatzenpichler, B. Bothner, T. R. McDermott, Aerobic bacterial methane synthesis. Proc Natl Acad Sci USA 118, e2019229118 (2021).

20. M. Bižić, T. Klintzsch, D. Ionescu, M. Y. Hindiyeh, M. Günthel, A. M. Muro-Pastor, W. Eckert, T. Urich, F. Keppler, H.-P. Grossart, Aquatic and terrestrial cyanobacteria produce methane. Science Advances 6, eaax5343 (2020).

21. T. Klintzsch, G. Langer, G. Nehrke, A. Wieland, K. Lenhart, F. Keppler, Methane production by three widespread marine phytoplankton species: release rates, precursor compounds, and potential relevance for the environment. Biogeosciences 16, 4129–4144 (2019).

22. Y. Mao, T. Lin, H. Li, R. He, K. Ye, W. Yu, Q. He, Aerobic methane production by phytoplankton as an important methane source of aquatic ecosystems: Reconsidering the global methane budget. Science of The Total Environment 907, 167864 (2024).

23. F. Janssen, M. Huettel, U. Witte, Pore-water advection and solute fluxes in permeable marine sediments (II): Benthic respiration at three sandy sites with different permeabilities (German Bight, North Sea). Limnology and Oceanography 50, 779–792 (2005).

24. M. Huettel, P. Berg, J. E. Kostka, Benthic Exchange and Biogeochemical Cycling in Permeable Sediments. Annu. Rev. Mar. Sci. 6, 23–51 (2014).

25. Y.-J. Chen, P. M. Leung, J. L. Wood, S. K. Bay, P. Hugenholtz, A. J. Kessler, G. Shelley, D. W. Waite, A. E. Franks, P. L. M. Cook, C. Greening, Metabolic flexibility allows bacterial habitat generalists to become dominant in a frequently disturbed ecosystem. The ISME Journal 15, 2986–3004 (2021).

26. J. E. Cable, G. C. Bugna, W. C. Burnett, J. P. Chanton, Application of 222Rn and CH4 for assessment of groundwater discharge to the coastal ocean. Limnology and Oceanography 41, 1347–1353 (1996).

27. G. Kim, D.-W. Hwang, Tidal pumping of groundwater into the coastal ocean revealed from submarine 222Rn and CH4 monitoring. Geophysical Research Letters 29, 23-1-23–4 (2002).

28. R. Hedderich, W. B. Whitman, “Physiology and Biochemistry of the Methane-Producing Archaea” in The Prokaryotes: Volume 2: Ecophysiology and Biochemistry, M. Dworkin, S. Falkow, E. Rosenberg, K.-H. Schleifer, E. Stackebrandt, Eds. (Springer, New York, NY, 2006; 10.1007/0-387-30742-7_34), pp. 1050–1079.

29. R. P. Gunsalus, J. A. Romesser, R. S. Wolfe, Preparation of coenzyme M analogs and their activity in the methyl coenzyme M reductase system of Methanobacterium thermoautotrophicum. Biochemistry 17, 2374–2377 (1978).

30. T. M. Webster, A. L. Smith, R. R. Reddy, A. J. Pinto, K. F. Hayes, L. Raskin, Anaerobic microbial community response to methanogenic inhibitors 2-bromoethanesulfonate and propynoic acid. Microbiologyopen 5, 537–550 (2016).

31. D. M. Karl, L. Beversdorf, K. M. Björkman, M. J. Church, A. Martinez, E. F. Delong, Aerobic production of methane in the sea. Nature Geosci 1, 473–478 (2008).

32. D. J. Repeta, S. Ferrón, O. A. Sosa, C. G. Johnson, L. D. Repeta, M. Acker, E. F. DeLong, D. M. Karl, Marine methane paradox explained by bacterial degradation of dissolved organic matter. Nature Geosci 9, 884–887 (2016).

33. Y. Li, C. G. Fichot, L. Geng, M. G. Scarratt, H. Xie, The Contribution of Methane Photoproduction to the Oceanic Methane Paradox. Geophysical Research Letters 47, e2020GL088362 (2020).

34. K.-Q. Xiao, F. Beulig, H. Røy, B. B. Jørgensen, N. Risgaard-Petersen, Methylotrophic methanogenesis fuels cryptic methane cycling in marine surface sediment. Limnology and Oceanography 63, 1519–1527 (2018).

35. J. M. Kurth, H. J. M. Op den Camp, C. U. Welte, Several ways one goal-methanogenesis from unconventional substrates. Appl Microbiol Biotechnol 104, 6839–6854 (2020).

36. S. Schorn, S. Ahmerkamp, E. Bullock, M. Weber, C. Lott, M. Liebeke, G. Lavik, M. M. M. Kuypers, J. S. Graf, J. Milucka, Diverse methylotrophic methanogenic archaea cause high methane emissions from seagrass meadows. PNAS 119 (2022).

37. M. B. Burg, J. D. Ferraris, Intracellular Organic Osmolytes: Function and Regulation *. Journal of Biological Chemistry 283, 7309–7313 (2008).

38. D. C. Yoch, Dimethylsulfoniopropionate: Its Sources, Role in the Marine Food Web, and Biological Degradation to Dimethylsulfide. Appl Environ Microbiol 68, 5804–5815 (2002).

39. A. N. Roychoudhury, E. Viollier, P. Van Cappellen, A plug flow-through reactor for studying biogeochemical reactions in undisturbed aquatic sediments. Applied Geochemistry 13, 269–280 (1998).

40. C. Pallud, C. Meile, A. M. Laverman, J. Abell, P. Van Cappellen, The use of flow-through sediment reactors in biogeochemical kinetics: Methodology and examples of applications. Marine Chemistry 106, 256–271 (2007).

41. I. R. Santos, B. D. Eyre, M. Huettel, The driving forces of porewater and groundwater flow in permeable coastal sediments: A review. Estuarine, Coastal and Shelf Science 98, 1–15 (2012).

42. J. T. Shaw, G. Allen, P. Barker, J. R. Pitt, D. Pasternak, S. J.-B. Bauguitte, J. Lee, K. N. Bower, M. C. Daly, M. F. Lunt, A. L. Ganesan, A. R. Vaughan, F. Chibesakunda, M. Lambakasa, R. E. Fisher, J. L. France, D. Lowry, P. I. Palmer, S. Metzger, R. J. Parker, N. Gedney, P. Bateson, M. Cain, A. Lorente, T. Borsdorff, E. G. Nisbet, Large Methane Emission Fluxes Observed From Tropical Wetlands in Zambia. Global Biogeochemical Cycles 36, e2021GB007261 (2022).

43. C. Helfter, M. Gondwe, M. Murray-Hudson, A. Makati, U. Skiba, From sink to source: high inter-annual variability in the carbon budget of a Southern African wetland. Philosophical Transactions of the Royal Society A: Mathematical, Physical and Engineering Sciences 380, 20210148 (2021).

44. S. D. Bridgham, K. Updegraff, J. Pastor, Carbon, Nitrogen, and Phosphorus Mineralization in Northern Wetlands. Ecology 79, 1545–1561 (1998).

45. K. Updegraff, J. Pastor, S. D. Bridgham, C. A. Johnston, Environmental and Substrate Controls over Carbon and Nitrogen Mineralization in Northern Wetlands. Ecological Applications 5, 151–163 (1995).

46. D. W. Valentine, E. A. Holland, D. S. Schimel, Ecosystem and physiological controls over methane production in northern wetlands. Journal of Geophysical Research: Atmospheres 99, 1563–1571 (1994).

47. S. L. Tsola, Y. Zhu, Y. Chen, I. A. Sanders, C. K. Economou, V. Brüchert, Ö. Eyice, Methanolobus use unspecific methyltransferases to produce methane from dimethylsulphide in Baltic Sea sediments. Microbiome 12, 3 (2024).

48. H. Seedorf, A. Dreisbach, R. Hedderich, S. Shima, R. K. Thauer, F420H2 oxidase (FprA) from Methanobrevibacter arboriphilus, a coenzyme F420-dependent enzyme involved in O2 detoxification. Arch Microbiol 182, 126–137 (2004).

49. A. L. Brioukhanov, A. I. Netrusov, R. I. L. Eggen, The catalase and superoxide dismutase genes are transcriptionally up-regulated upon oxidative stress in the strictly anaerobic archaeon Methanosarcina barkeri. Microbiology 152, 1671–1677 (2006).

50. A. Das, E. D. Coulter, D. M. Kurtz, L. G. Ljungdahl, Five-Gene Cluster in Clostridium thermoaceticumConsisting of Two Divergent Operons Encoding Rubredoxin Oxidoreductase-Rubredoxin and Rubrerythrin–Type A Flavoprotein– High-Molecular-Weight Rubredoxin. Journal of Bacteriology 183, 1560–1567 (2001).

51. R. Sheehan, A. C. McCarver, C. E. Isom, E. A. Karr, D. J. Lessner, The Methanosarcina acetivorans thioredoxin system activates DNA binding of the redox-sensitive transcriptional regulator MsvR. J Ind Microbiol Biotechnol 42, 965–969 (2015).

52. D. Susanti, J. H. Wong, W. H. Vensel, U. Loganathan, R. DeSantis, R. A. Schmitz, M. Balsera, B. B. Buchanan, B. Mukhopadhyay, Thioredoxin targets fundamental processes in a methane-producing archaeon, Methanocaldococcus jannaschii. Proceedings of the National Academy of Sciences 111, 2608–2613 (2014).

53. Z. Lyu, Y. Lu, Metabolic shift at the class level sheds light on adaptation of methanogens to oxidative environments. ISME J 12, 411–423 (2018).

54. Y.-J. Chen, P. M. Leung, P. L. M. Cook, W. W. Wong, T. Hutchinson, V. Eate, A. J. Kessler, C. Greening, Hydrodynamic disturbance controls microbial community assembly and biogeochemical processes in coastal sediments. ISME J (2021b).

55. R. Wilms, H. Sass, B. Köpke, H. Cypionka, B. Engelen, Methane and sulfate profiles within the subsurface of a tidal flat are reflected by the distribution of sulfate-reducing bacteria and methanogenic archaea. FEMS Microbiology Ecology 59, 611–621 (2007).

56. W. S. Reeburgh, Oceanic methane biogeochemistry. Chemical Reviews 107, 486–513 (2007).

57. E. Precht, U. Franke, L. Polerecky, M. Huettel, Oxygen dynamics in permeable sediments with wave-driven pore water exchange. Limnology and Oceanography 49, 693–705 (2004).

58. M. Huettel, W. Ziebis, S. Forster, G. W. Luther, Advective Transport Affecting Metal and Nutrient Distributions and Interfacial Fluxes in Permeable Sediments. Geochimica et Cosmochimica Acta 62, 613–631 (1998).

59. A. V. Borges, W. Champenois, N. Gypens, B. Delille, J. Harlay, Massive marine methane emissions from near-shore shallow coastal areas. Sci Rep 6, 27908 (2016).

60. N. Gypens, A. V. Borges, Increase in dimethylsulfide (DMS) emissions due to eutrophication of coastal waters offsets their reduction due to ocean acidification. Frontiers in Marine Science 1 (2014).

61. A. Oschlies, A committed fourfold increase in ocean oxygen loss. Nat Commun 12, 2307 (2021).

62. Q. Xing, L. Tosi, F. Braga, X. Gao, M. Gao, Interpreting the progressive eutrophication behind the world’s largest macroalgal blooms with water quality and ocean color data. Nat Hazards 78, 7–21 (2015).

63. L. A. Green-Gavrielidis, C. S. Thornber, Will Climate Change Enhance Algal Blooms? The Individual and Interactive Effects of Temperature and Rain on the Macroalgae Ulva. Estuaries and Coasts 45, 1688–1700 (2022).

64. R. F. Weiss, B. A. Price, Nitrous oxide solubility in water and seawater. Marine Chemistry 8, 347–359 (1980).

65. D. A. Wiesenburg, N. L. Jr. Guinasso, Equilibrium solubilities of methane, carbon monoxide, and hydrogen in water and sea water. J. Chem. Eng. Data 24, 356–360 (1979).

66. B. Buchfink, C. Xie, D. H. Huson, Fast and sensitive protein alignment using DIAMOND. Nat Methods 12, 59–60 (2015).

67. H. R. Gruber-Vodicka, B. K. B. Seah, E. Pruesse, phyloFlash: Rapid Small-Subunit rRNA Profiling and Targeted Assembly from Metagenomes. mSystems 5, 10.1128/msystems.00920-20 (2020).

68. W. E. Wentworth, S. V. Vasnin, S. D. Stearns, C. J. Meyer, Pulsed discharge helium ionization detector. Chromatographia 34, 219–225 (1992).

## References in Supplementary only

64. V. W. Salazar, B. Shaban, M. del M. Quiroga, R. Turnbull, E. Tescari, V. Rossetto Marcelino, H. Verbruggen, K.-A. Lê Cao, Metaphor—A workflow for streamlined assembly and binning of metagenomes. GigaScience 12, giad055 (2023).

65. S. Chen, Ultrafast one-pass FASTQ data preprocessing, quality control, and deduplication using fastp. iMeta 2, e107 (2023).

66. D. Li, C.-M. Liu, R. Luo, K. Sadakane, T.-W. Lam, MEGAHIT: an ultra-fast single-node solution for large and complex metagenomics assembly via succinct de Bruijn graph. Bioinformatics 31, 1674–1676 (2015).

67. D. Hyatt, G.-L. Chen, P. F. LoCascio, M. L. Land, F. W. Larimer, L. J. Hauser, Prodigal: prokaryotic gene recognition and translation initiation site identification. BMC Bioinformatics 11, 119 (2010).

68. S. F. Altschul, T. L. Madden, A. A. Schäffer, J. Zhang, Z. Zhang, W. Miller, D. J. Lipman, Gapped BLAST and PSI-BLAST: a new generation of protein database search programs. Nucleic Acids Research 25, 3389–3402 (1997).

69. P.-A. Chaumeil, A. J. Mussig, P. Hugenholtz, D. H. Parks, GTDB-Tk v2: memory friendly classification with the genome taxonomy database. Bioinformatics 38, 5315–5316 (2022).

70. R. C. Edgar, Muscle5: High-accuracy alignment ensembles enable unbiased assessments of sequence homology and phylogeny. Nat Commun 13, 6968 (2022).

71. S. Kalyaanamoorthy, B. Q. Minh, T. K. F. Wong, A. von Haeseler, L. S. Jermiin, ModelFinder: fast model selection for accurate phylogenetic estimates. Nat Methods 14, 587–589 (2017).

72. L.-T. Nguyen, H. A. Schmidt, A. von Haeseler, B. Q. Minh, IQ-TREE: A Fast and Effective Stochastic Algorithm for Estimating Maximum-Likelihood Phylogenies. Molecular Biology and Evolution 32, 268–274 (2015).

73. I. Letunic, P. Bork, Interactive Tree of Life (iTOL) v6: recent updates to the phylogenetic tree display and annotation tool. Nucleic Acids Res 52, W78–W82 (2024).

74. N. US Department of Commerce, Global Monitoring Laboratory - Carbon Cycle Greenhouse Gases. https://gml.noaa.gov/ccgg/trends_ch4/.

75. R. F. Weiss, B. A. Price, Nitrous oxide solubility in water and seawater. Marine Chemistry 8, 347–359 (1980).

76. D. A. Wiesenburg, N. L. Jr. Guinasso, Equilibrium solubilities of methane, carbon monoxide, and hydrogen in water and sea water. J. Chem. Eng. Data 24, 356–360 (1979).

77. B. Buchfink, C. Xie, D. H. Huson, Fast and sensitive protein alignment using DIAMOND. Nat Methods 12, 59–60 (2015).

78. H. R. Gruber-Vodicka, B. K. B. Seah, E. Pruesse, phyloFlash: Rapid Small-Subunit rRNA Profiling and Targeted Assembly from Metagenomes. mSystems 5, 10.1128/msystems.00920-20 (2020).

79. W. E. Wentworth, S. V. Vasnin, S. D. Stearns, C. J. Meyer, Pulsed discharge helium ionization detector. Chromatographia 34, 219–225 (1992).

