## Supplementary files for "Aerotolerant methanogens use seaweed and seagrass metabolites to drive marine methane emissions"

**The file includes:**

Materials and Methods

Fig. S1 – S7

Tables S1 – S2

**Other Supplementary Materials for this manuscript include the following:**

Scripts accessible at https://github.com/GreeningLab/Sand-methanogen-manuscript

Sequence data available at NCBI Sequence Read Archive Accession: PRJNA1165813

### Materials and Methods

#### Field measurements and sampling sites

*In situ* measurements and sediment sampling were conducted in Australia at Werribee, St Kilda and Shoreham. In Denmark, samples were collected from Avernakø. For coordinates and dates of sampling, see Table S1. These sites were selected for their different macrophyte accumulation characteristics. For details of each site, see Fig. S7 (photos of each site) and Tables S1.1-S1.2 (descriptions, coordinates, and measured methane concentrations).

Seawater samples were collected for *in situ* dissolved methane ($CH_4$) analysis, and were collected with minimal gas loss by over-filling 12 mL exetainers from the bottom up using syringes fit with 10 cm low gas-permeability tubing (MasterFlex 06401-16). Samples were preserved by adding 50 µL saturated $HgCl_2$ and then stored at room temperature. For dissolved methane concentration, samples were analyzed by GC PDHID (gas chromatography interfaced with pulsed-discharge helium ionization detector). For analysis details see chemical analysis section below.

#### Flow through reactors (FTRs)

Flow through reactors were chosen as a controlled incubation setup that allowed advection through sediment at rates similar to *in situ* conditions, as well as allowing time-resolved rate measurements of production or consumption of methane and other compounds such as dissolved inorganic carbon and oxygen.

Sand was collected using acrylic cores from the shallow subtidal zone and transported back to the lab. The top 0-5 cm were sectioned then pooled, sieved (1 mm) to remove shells and other debris, and homogenized in locally -collected fully oxygenated seawater. Sand was then packed into FTRs (diameter 48 mm, length 20 mm) underwater and inlet connected to a reservoir (Fig. S1). Reservoirs contained seawater collected from the same site and were bubbled continuously with $N_2$ + $CO_2$ (820 ppm) to remove oxygen while maintaining pH (anoxic treatments) or lab air (oxic treatments). Control reservoirs contained only seawater while treatment reservoirs contained seawater + macrophyte extract (20:1 ratio, chosen as an attempt to simulate a large macrophyte accumulation event) or seawater with 10 µM methylphosphonate (MPn). Seawater was pumped through FTRs using a peristaltic pump set to 45 mL/h and not recirculated. Reservoirs were topped up with purged seawater approximately every 24 hours and the macrophyte extract ratio or MPn concentration maintained.

To enhance reproducibility across FTR experiments from different sites, a macrophyte extract was prepared as a consistent source of dissolved macrophyte metabolites. We collected mixed macrophytes from Shoreham, consisting of approximately 60% seagrass (*Amphibolis antarctica*) and the remainder as mixed seaweeds. A total of 278 g (dry weight) of mixed macrophytes was combined with 9 L of seawater and stored at 4 °C for nine days. Solids were then strained out and the liquid extract was frozen until use.

Dissolved oxygen and pH in the reservoirs and FTR outlets were monitored with optical flow through-cell oxygen sensors (Firesting Oxygen meter, Pyroscience) and Hach HQ40d pH meter, respectively.

Samples for $CH_4$ and dissolved inorganic carbon (DIC) measurements were taken from the reservoir and FTR outlet using gas-tight glass syringes, and processed as for field samples.

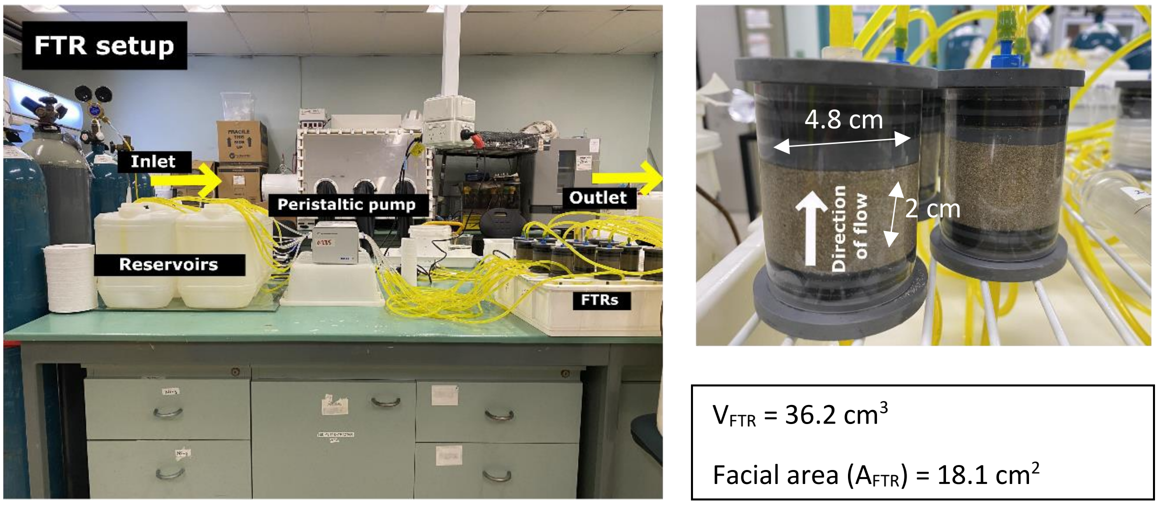


Figure S1. Diagram of lab flow through reactor (FTR) setup (left) and close up of FTR (right) including dimensions of sand cavity.

Calculations to normalize FTR methane production rates to wetland flux rates were performed as follows. We chose 0.5 cm as the sediment depth over which to integrate the methane production rate (rather than 2 cm as was the length of our FTRs) as this is a more conservative estimate of the advective penetration of high-substrate surface water under wave-pumping in the intertidal zone (*41*).

$$Maximum\ rate\ of\ methane\ production = x \approx 48\ \mu mol \cdot cm_{sand}^{-3} \cdot h^{-1}$$

$$Surface\ area\ molar\ flux\ rate\ per\ FTR\ integrated\ over\ 0.5\ cm$$

$$= x * \frac{V_{FTR}}{4} / A_{FTR}$$

$$= 24.0\ \mu mol \cdot cm^{-2} \cdot h^{-1}$$

$$Unit\ conversion\ to\ mmol\ per\ m^2 = 240\ mmol \cdot m^{-2} \cdot h^{-1}$$

$$Mass\ flux\ rate\ per\ m^2 = 240 * \frac{16.04}{1000} = 3.8\ g \cdot m^{-2} \cdot h^{-1}$$

Slurries

Sand was collected and processed as described for FTRs. For each sediment slurry, 30 g sediment was transferred into a 160 mL serum vial by flushing down a funnel with 70 mL seawater collected from the same location as the sediment. For seawater-only and seawater + macrophyte slurries, 100 mL seawater was used, with or without 3 g wet weight of macrophyte. All treatments consisted of triplicate slurries.

Vials were crimp-closed with butyl rubber stoppers and the headspace was purged for 2 min with $N_2$, shaken for 2 min, then purged again to remove oxygen. To sample for $CH_4$ measurement, 2 mL He was injected into the headspace and gently agitated before a 2 mL sample was taken from the headspace and injected into a He-purged 3 mL exetainer. The samples were analyzed by GC PDHID (see chemical analysis).

To examine the relative importance of sediment and macrophyte in methane production, slurries were prepared in combinations of seawater, sediment, and macrophyte. Macrophytes were either brown drift algae collected from Werribee (Fig. 1D) or seagrass collected from Shoreham (*Amphibolis antarctica*) (Fig. S6).

To examine inhibition of methanogens, sediment slurries were prepared with or without 20 mM BES (2-bromoethane sulfonate) in the presence or absence of spirulina (dried, powdered *Arthrospira platensis* sourced from Nature's Way®, 12.5 mg per slurry).

To identify the effect of different substrates on $CH_4$ production pathways, trimethylamine (TMA), dimethylsulfide (DMS), methylamine, choline, methylphosphonate (MPn) or acetate was added to separate slurries. All chemicals were sourced from Sigma Aldrich and the final concentration of the substrate in the slurries was adjusted to 100 µM. For hydrogenotrophic pathway, hydrogen gas ($H_2$) was added directly to the headspace and equilibrated to a final concentration of 340 ± 20 ppm.

Time 0 was defined as the point when oxygen was completely purged from the slurries and the substrates added, after which an initial sample was immediately taken. The incubations were run for at least 50 and up to 400 hours depending on how quickly methane was produced (i.e. they were incubated until a distinct increase in methane concentration was detected in at least one treatment) with samples taken approximately every 24 hours.

### Chemical analysis

#### *Analysis of $CH_4$ by gas chromatography (GC)*

For analysis of dissolved methane samples (all *in situ* and FTR samples), a headspace was introduced to the 12 mL exetainer by replacing 5 mL of seawater sample with high purity Helium gas. Samples were then shaken vigorously for 4 min, and gas in the headspace was analyzed using a gas chromatograph interfaced with a pulsed discharge helium ionization detector (GC-PDHID, Valco Instruments Co. Inc.) (*52*). The original dissolved gas concentration was calculated according to Wiesenburg and Guinasso (1979) and Weiss and Price (1980) (*53, 54*). Slurry headspace samples were injected directly.

For the pure culture experiments the methane concentration was too high for GC PDHID and so were analyzed by GC FID (flame ionization detector).

100 µL samples were taken directly from the headspace with an SGE gas-tight syringe and injected manually into a PerkinElmer Clarus® 580 gas chromatograph fitted with an SGE BP20 wax column (30 m length, 0.32 mm diameter, 1 µm film thickness).

Manual triplicate five-point calibration was performed at the start of each run along with standards and blanks at the end of each run. Calibration gases used were NATA-accredited calibration gases from Air Liquide and BOC HiQ.

#### *Analysis of dissolved inorganic carbon (DIC)*

For analysis of DIC, a 1 mL seawater sample from FTRs and slurries was acidified using 1M $H_3PO_4$ to convert carbonate and bicarbonate to carbon dioxide. The carbon dioxide produced was stripped from the liquid phase by bubbling with ultrapure nitrogen gas and analyzed using a LI-COR non-dispersive infrared analyzer (Apollo SciTech). Standard reference material for three-point calibration was obtained from Scripps Institute of Oceanography, University of California, San Diego.

#### *Radon analysis for groundwater input*

Radon ($^{222}Rn$) concentration was analyzed as a tracer for groundwater using $^{222}Rn$ in-air monitor (RAD7; Durridge Company). 1L of water samples was collected in Schott bottles and capped underwater where possible to eliminate bubbles. The samples were analyzed in the lab within 24 hours of collection. For analysis, 500 mL sample was degassed for 5 min into a closed-air loop of known volume and was counted for 2 hours. The concentrations were corrected for salinity, temperature and decay time loss to accurately represent *in situ* concentrations.

### Metagenomics

Samples for metagenomic analysis were subsampled from sediment cores collected for FTR experiments (0-5 cm depth). Additional samples for metagenomic analysis were subsampled from Shoreham FTR sediment at the end of the experiment (T = ~60 h).

Total community DNA was extracted from 0.25 g of sediment using QIAGEN DNeasy PowerSoil Pro extraction kit according to manufacturer's instructions. The DNA yield was measured by a Qubit 2.0 Fluorometer (Invitrogen). Shotgun sequencing was conducted by Micromon Genomics using DNBSEQ at 10 Gb depth. The Metaphor pipeline (*64*) was employed for read quality control, assembly, and binning. Specifically, raw reads from the eight metagenome libraries underwent quality control by trimming primers and adapters, removal of artifacts and low-quality reads using fastp (*65*) with parameters length_required: 50, cut_mean_quality: 30, and extra: --detect_adapter_for_pe. Kept reads were co-assembled using MEGAHIT v1.2.9 (*66*) with default settings. Contigs shorter than 1,000 bp were discarded. Open reading frames (ORFs) were predicted using Prodigal v2.6.2.9 (*67*), then annotated using homology-based searches and hidden Markov model (HMM) searches using DIAMOND blastp (*68*) against a custom protein database of 51 metabolic marker genes as described (https://bridges.monash.edu/collections/_/5230745/3). Full scripts and settings are provided in https://github.com/GreeningLab/Sand-methanogen-manuscript

### Methanogen isolation and whole genome sequencing

Isolation experiments were undertaken from the surface 5 cm of sediment of two highly macrophyte-impacted sandy beaches on Avernakø and Shoreham, collected on 10/07/2023 and 22/01/2024 respectively. Methylotrophic methanogens were enriched in slurries prepared as above, with addition of 1 mM trimethylamine chloride (TMA) and incubation at 15˚C˚ (Denmark) or 19-21˚C21˚ (Australia) for four weeks. A methanogen was then isolated from each enrichment by three rounds of dilution to extinction in modified DSMZ 141c *Methanococcoides* medium (modifications were addition of 0.5 mg/L sodium resazurin, ampicillin 100 mg/L and kanamycin 200 mg/L, and exclusion of yeast extract, sodium acetate and trypticase peptone). Purity of the isolate was ensured by adding antibiotics to a methanogen-selective isolation media. The growth of the methanogenic strains was monitored by $OD_{600}$ measurements, optical microscopy, and methane production and the resulting strains referred to as DA (Denmark) and SH (Australia). Purity was then confirmed through epifluorescence microscopy, targeting the F420-autofluorescence typical of methanogens, along with PCR screening (DA only) and sequencing (both strains). For whole genome sequencing (WGS), DNA was extracted using the DNeasy PowerSoil Pro kit according to the manufacturer's instructions. For strain DA, DNA concentration and purity were measured with the Qubit dsDNA HS Assay kit (Thermo Fisher Scientific) and NanoDrop One, respectively. DNA size distributions were evaluated using the Genomic DNA ScreenTapes on Agilent Tapestation 4200. A barcoded SQK-NBD114.95 DNA library was prepared (Oxford Nanopore Technologies) and loaded onto a primed FLO-PRO114M (R10.4.1) flow cell and sequenced on a PromethION P24 device.

For strain SH, next generation sequencing library preparations were constructed following the manufacturer's protocol (VAHTS Universal Plus DNA Library Prep Kit for Illumina V2). Libraries with different indexes were multiplexed and loaded on an Illumina NovaSeq 6000 instrument according to manufacturer's instructions (Illumina, San Diego). Sequencing was carried out using a 2x150 paired-end (PE) configuration; image analysis and base calling were conducted by the NovaSeq Control Software (NVCS) + RTA 3 (Illumina) on the NovaSeq 6000 instrument.

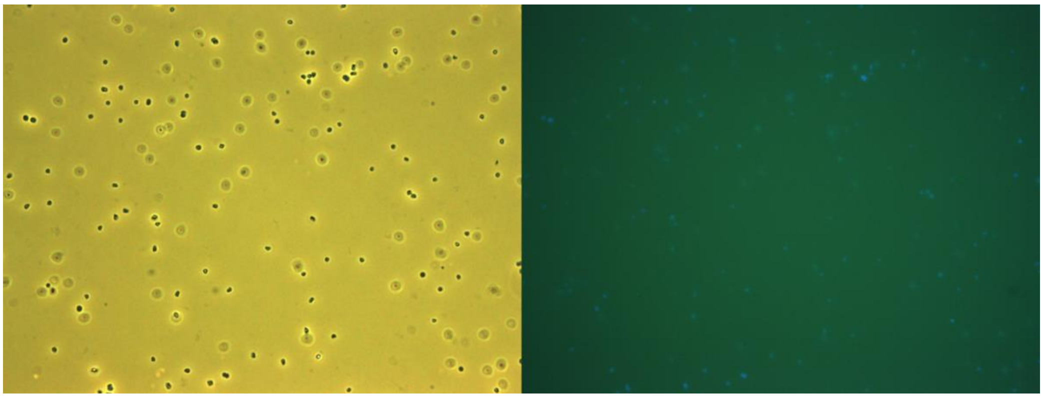

Figure S2. Visible light and epifluorescence microscope images of DA.

<u>Phylogenetic tree building and visualization</u>

We used GTDB-tk (*69*) to align the genomes of the two isolates with 71 reference methanogens genomes, whereas we used Muscle (*70*) to align the three *mcrA* contigs with 196 and 31 *mcrA* reference sequences for producing the alignments used to generate the supplementary and main *mcrA* phylogenetic trees, respectively. Phylogenetic trees were generated using IQ-TREE v2.2.2.6 (*71*, *72*) and 1,000 ultrafast bootstraps. Genome tree was generated with model LG+F+I+R6, *mcrA* supplementary tree alignment was generated with model LG+F+I+R5; and *mcrA* main tree alignment was generated with model LG+I+G4. We plotted the phylogenetic trees with genome statistics using iTOL v6 (*73*). Full scripts and settings are provided in https://github.com/GreeningLab/Sand-methanogen-manuscript

<u>Oxygen pulse exposure experiment</u>

Isolate cultures were grown on TMA to stationary phase in 20 mL crimp-top vials as for isolation but with contactless autoclavable oxygen sensor spots inserted at the bottom (PyroScience OXSP5 oxygen sensor spots, calibrated according to manufacturer's instructions). At the beginning of the experiment, approximately 10 mL of lab air was injected into the headspace of treatment vials, with an outlet needle inserted to prevent overpressure, and the vials shaken to equilibrium. Oxygen readings were taken every 5 seconds in all oxygen exposure treatment vials plus one no oxygen exposure vial (as the maximum number of channels measuring oxygen concurrently is four). After 30 minutes all vials were flushed with helium to remove oxygen and residual methane. As soon as the dissolved oxygen readings in oxygen exposure treatments were stable at <0.01 mg/L and no pink resazurin was visible, 100 µL of sterile anoxic 500 g/L TMA was then injected into all vials (including the media control) and methane measurements began. Methane in the headspace was analyzed approximately every hour for 5 hours. For strain SH, control and treatment were prepared in duplicate, and strain DA in triplicate.

**Supplementary Figures**

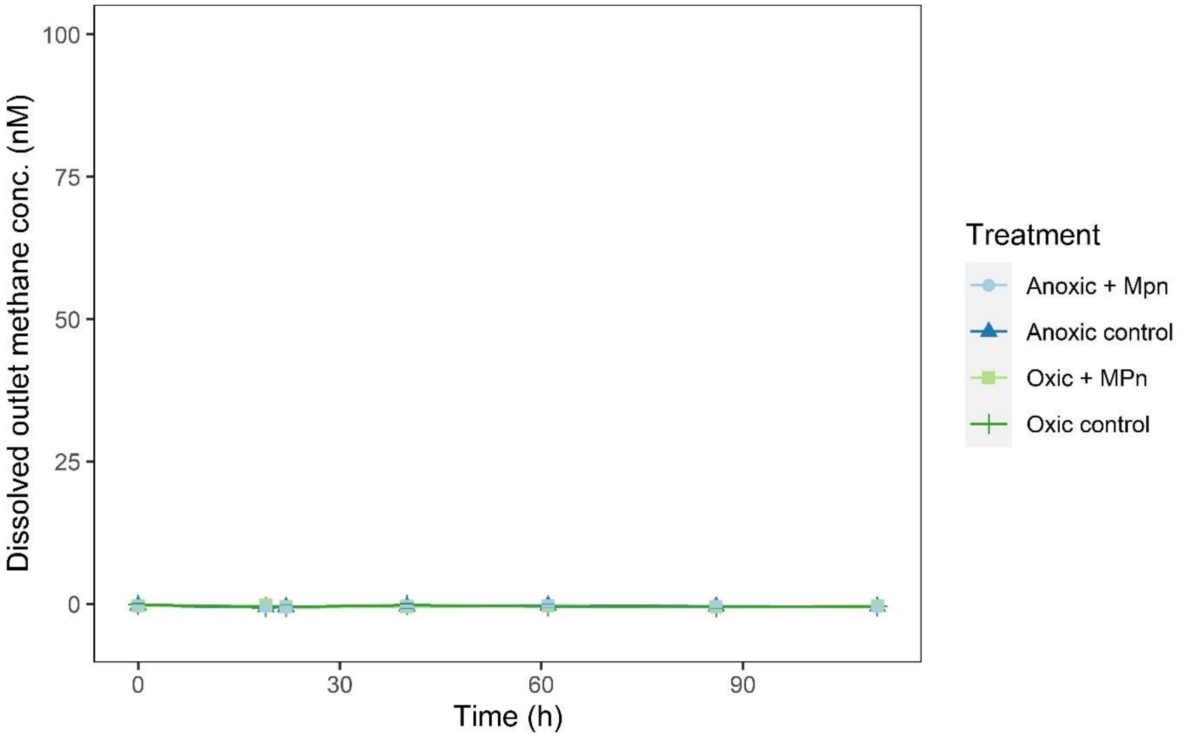


Figure S3. Oxic and anoxic FTRs (surface 0-5 cm Werribee sediments collected 14/10/2021) with and without addition of methylphosphonate (10 µM) in seawater reservoir.

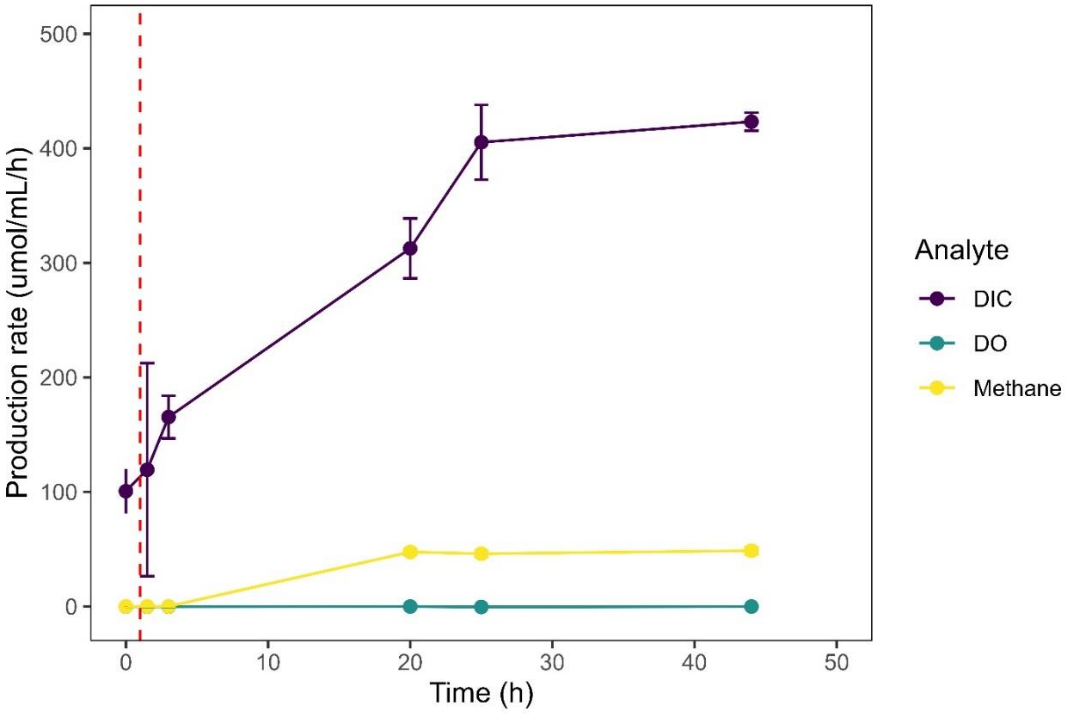


Figure S4. Methane, dissolved oxygen (DO) and dissolved inorganic carbon production rate in Shoreham FTRs with macrophyte extract, normalized to $\mu mol/mL_{sediment}/h$.

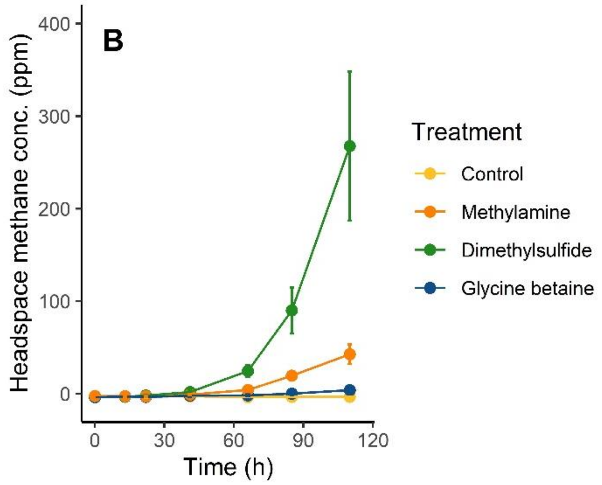


Figure S5. Sediment slurry experiments (0-5 cm depth at Werribee) with targeted substrate addition (100 µM final concentration for all substrates) sediment collected 14/10/2021. Error bars represent one standard deviation from triplicate slurries.

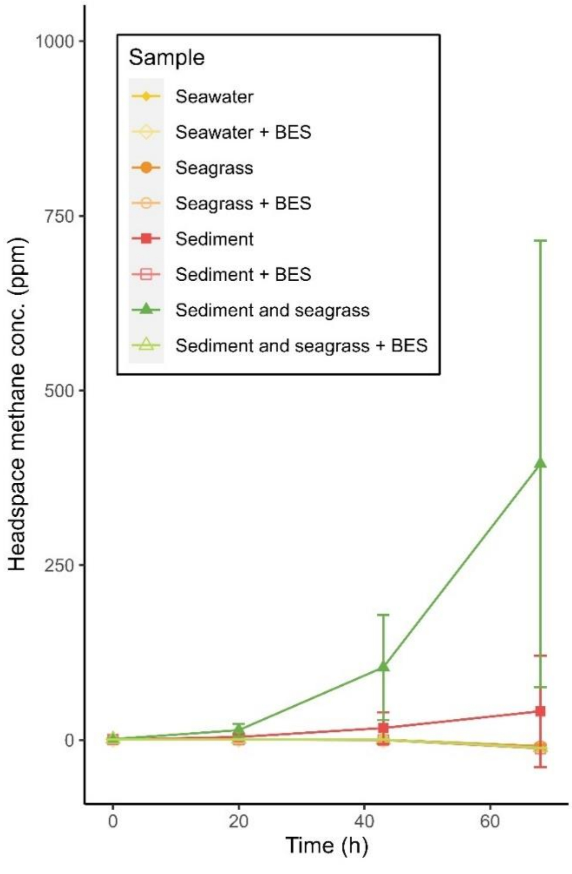


Fig S6. Slurry experiments with combinations of seawater, seagrass, and surface sediment (0-5 cm depth at Shoreham) as well as specific archaeal methanogenesis inhibitor 2-bromoethane sulfonate (BES) 20 mM.  Error bars represent one standard deviation from triplicate slurries.

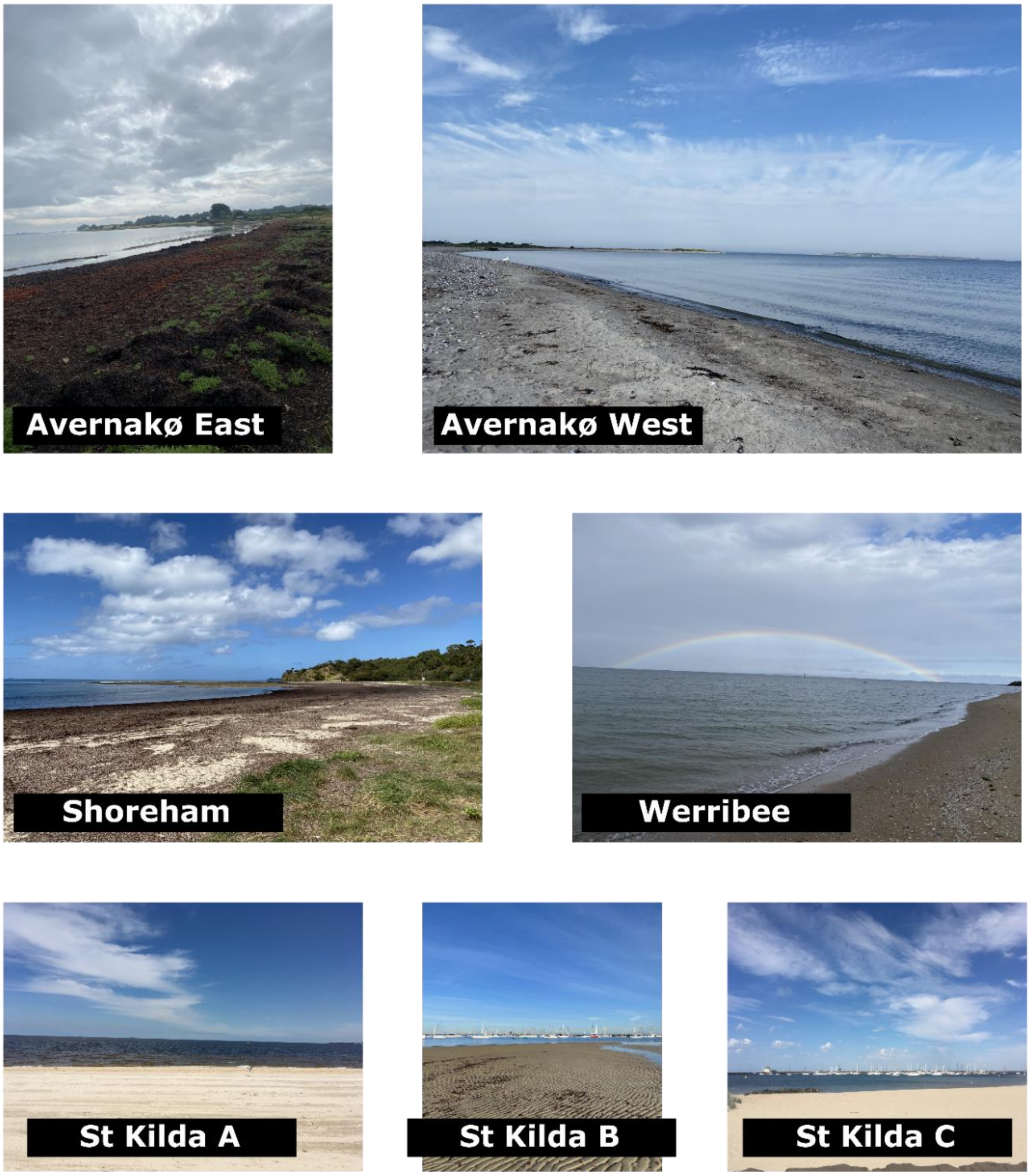


Figure S7. Photos of all sampling sites in the study.

Table S1. Field site descriptions and macrophyte accumulation characteristics.

| Site | Description and macrophyte accumulation characteristics |
|---|---|
| **Avernakø** | Two beaches on East (Av 1, 2, 3) and West (Av 4, 5) sides of Avernakø boat harbor. Avernakø East had large accumulations of macrophyte (very degraded so identification was not possible) while Avernakø West was almost bare, with small isolated pieces of ulva |
| **St Kilda A** | Exposed high energy beach (relative to other St Kilda sites) with no visible macrophyte accumulation. |
| **St Kilda B** | Sandy beach at entrance to boat harbor and partially protected by a breakwater. Contains very. Very small isolated pieces of seaweed (mostly ulva) |
| **St Kilda C** | Sandy beach in boat harbor and highly protected by a breakwater. Contains very. Very small isolated pieces of seaweed (mostly ulva) |
| **Shoreham** | Large mounds of macrophyte biomass partially submerged in surf. Approximately 60% seagrass (primarily *Amphibolis antarctica*) with remainder mixed seaweeds including ulva, kelp, and filamentous red species. |
| **Werribee** | A history of large seasonal deposits of drift algae (brown). Radon/methane correlation study conducted at the tail end of one of these blooms with one small remaining patch. All other experiments (FTRs and slurries) conducted when no seaweed was present. |

Table S2. Field site locations, average surface water methane concentrations (n=3), and calculated percent saturation. Av = Avernakø, Denmark. SK = St Kilda (A-C), Sh = Shoreham, WSB = Werribee Southern Beach, Australia. Numbers refer to subsites, usually approx. 20 meters apart forming a transect along the beach. Calculated % saturation uses equilibrium concentration calculated using 1922.39 ppb average annual atmospheric methane concentration (*74*).

| Site | Date sampled (DD/MM/YY) | Lat | Long | Average $CH_4$ conc. (nM) | Standard deviation | Calculated %saturation |
|---|---|---|---|---|---|---|
| Av1 | 22/05/23 | 55.03896 | 10.25353 | 2100 | 300 | 78156 |
| Av2 | 22/05/23 | 55.03853 | 10.2535 | 4200 | 200 | 155442 |
| Av3 | 22/05/23 | 55.03799 | 10.25357 | 4400 | 200 | 163706 |
| Av4 | 22/05/23 | 55.03978 | 10.25001 | 110 | 30 | 4088 |
| Av5 | 22/05/23 | 55.03954 | 10.24972 | 43 | 9 | 1586 |
| SKA1 | 06/10/22 | -37.8516 | 144.9553 | 10.1 | 0.9 | 373 |
| SKA2 | 06/10/22 | -37.8515 | 144.955 | 9.0 | 0.7 | 332 |
| SKA3 | 06/10/22 | -37.8514 | 144.9546 | 9.6 | 1.4 | 354 |
| SKB1 | 06/10/22 | -37.8611 | 144.9662 | 144 | 5 | 5350 |
| SKB2 | 06/10/22 | -37.8605 | 144.9655 | 106.7 | 0.9 | 3951 |
| SKB3 | 06/10/22 | -37.8605 | 144.9655 | 278 | 5 | 10282 |
| SKC1 | 06/10/22 | -37.8619 | 144.9699 | 66.5 | 0.9 | 2462 |
| SKC2 | 06/10/22 | -37.8616 | 144.9694 | 82.8 | 1.4 | 3068 |
| SKC3 | 06/10/22 | -37.8621 | 144.9693 | 17.0 | 0.3 | 628 |
| Sh1 | 13/09/22 | -38.4343 | 145.0477 | 200.9 | 0.4 | 7442 |
| Sh2 | 13/09/22 | -38.4349 | 145.0476 | 226 | 7 | 8365 |
| Sh3 | 13/09/22 | -38.435 | 145.0473 | 114 | 3 | 4223 |
| Sh4 | 13/09/22 | -38.4352 | 145.047 | 67.7 | 0.6 | 2499 |
| Sh5 | 13/09/22 | -38.4353 | 145.0468 | 119 | 13 | 4397 |

| | | | | | | |
|---|---|---|---|---|---|---|
| WSB 1 | 11/02/22 | -37.9705 | 144.7044 | 29.0 | 1.1 | 1075 |
| WSB 2 | 11/02/22 | -37.9707 | 144.7041 | 23.8 | 1.4 | 881 |
| WSB 3 | 11/02/22 | -37.9708 | 144.7038 | 24.4 | 0.5 | 902 |
| WSB 4 | 11/02/22 | -37.9711 | 144.7034 | 32.7 | 0.8 | 1210 |
| WSB 5 | 11/02/22 | -37.9713 | 144.7029 | 60 | 2 | 2236 |
